# Astrocyte Autophagy Response Upon Neuronal Cilia Loss in the Aging Brain

**DOI:** 10.1101/2022.06.14.496086

**Authors:** Laura de las Heras-García, Olatz Pampliega

**Affiliations:** Departamento de Neurociencias, Universidad del País Vasco (UPV/EHU), and Achucarro Basque Center for Neurosciences, 48940 – Leioa, Spain

**Keywords:** autophagy, primary cilium, astrocyte

## Abstract

Primary cilia are microtubule-based signaling organelles present in the plasma membrane of most cell types, including mature astrocytes and neurons. However, little is known about the role of this organelle in the mature brain. Data from our lab show that neuronal primary cilia (nPC) is required for soluble amyloid beta oligomer signaling and modulation of autophagy, and that these events are age dependent. Here, we hypothesize that astrocytes react to the loss of nPC and that aging might impact these events. For that purpose, we have characterized morphological changes in astrocytes as well as in the cilium and autophagy of these cells in brain tissue from young and old mice with impaired PC in neurons. Our results show that upon loss of PC in neurons astrocytes become reactive and reduce their lysosomal capacity, an effect that is reinforced with aging. Moreover, aging reduced the pool of ciliated astrocytes, which might impact their ability to react to extracellular events. Overall, our data suggest that the PC might act an intermediary in the communication between astrocytes and neurons.

**Highlights of the paper:**

- Astrocytes become reactive upon loss of primary cilia in neurons, which is reinforced during aging.
- Astrocytes in the old brain are less ciliated.
- Loss of neuronal primary cilia decreases lysosomal capacity in astrocytes in age-dependent manner.

## INTRODUCTION

Autophagy degrades cytoplasmic components including proteins and organelles in the lysosome. As its activity declines with age [1], malfunction of autophagy is a major component of age-dependent neurodegenerative diseases (NDD) like Alzheimer’s disease (AD), Parkinson’s disease (PD), or Huntington’s disease (HD). Loss of autophagy in neurons results in the intracellular accumulation of protein aggregates characteristic of the major NDD, leading in the long run to neuronal cell loss [2; 3]. Besides neurons, astrocytes are the most abundant cells in the brain, cells that are gaining relevance in the pathophysiology of aging and NDD [4; 5; 6]. However, autophagy in astrocytes has been poorly characterized, and therefore there is minimum knowledge about the role of astrocytic autophagy in brain diseases and aging.

Astrocytes and neurons are in constant communication to ensure a proper brain activity. Neuron-astrocyte intercellular communication is achieved via release of gliotransmmiters [7] or through extracellular vesicles [8]. However how these intercellular communication affects autophagy in the neighbor cell is not well understood.

Primary cilia (PC) are signaling organelles sitting on the surface of most cell types, including mature neurons and astrocytes [9]. Although the precise role of primary cilia in the adult brain remains unknown, increasing evidence point to a key role of this organelle in brain diseases, including NDD, mood disorders or aging [10; 11; 12]. Signaling from PC modulates starvation-induced autophagy [13], and recent work from our laboratory describes that extracellular amyloid beta oligomers require the presence of neuronal PC to impair autophagy function in an age-dependent manner [14].

Here we study whether astrocytes modify their morphology and autophagy upon loss of neuronal PC and aging. We hypothesize that astrocytes and neurons take advantage of the PC sensing role to detect changes in the neighbor cell types and modify their intracellular autophagy accordingly.

## MATERIALS AND METHODS

### Animals

For the purpose of this project, we used Ift88F/F and Ift88-SLICK-H (Iftt88F/F::Thy1-Cre/+) mice (hereafter referred to as F/F and Ift88-/-^Thy1^, respectively), using both sexes of mice at 3 and 15 months of age. Mice were maintained under standard conditions with food and water ad libitum in a normal light/dark cycle. All experimental procedures were approved by the Committee on Animal Health and Care of the University of Bordeaux and French Ministry of Agriculture and Forestry (authorization numbers, APAFIS#8430-2017010412591287 and APAFIS#8719-2017012411024773). Maximal efforts were made to reduce the suffering and the number of animals used.

PC loss (Ift88 deletion) in Thy1+-EYFP neurons was induced in CRE animals after intraperitoneal injection of Tamoxifen (75mg/kg during 5 consecutive days). Recombination was checked by PCR [14].

### Tissue preparation

Following sacrifice, mice were perfused with 4% paraformaldehyde in 0.1M phosphate buffer (PB). Brains (cortex, hippocampus, striatum, olfactory bulb) were dissected under microscope, dehydrated in 20% sucrose for 24h, frozen at -70ºC for 2min and preserved in OCT at -80ºC. 20µm coronal sections were cut using Leyca cryostat. Free-floating sections were collected and stored in PBS at 4ºC for a maximum of 7 days until passed to cryopreservation solution (30% glycerol, 30% ethylanglycol, 30% ddH2O, and PB 0.4M) and kept at -20ºC.

### Tissue immunohistochemistry

Free-floating brain slices (n = 3 per experimental condition) were washed three times 5 min in PBS and blocked in PBS containing 0.1% triton X-100, 5% NGS for 1h at RT. Thereafter, sections were incubated with primary antibody diluted in PBS containing 0.1% tween X-20, 5% NGS, ON at 4ºC rocking. Primary antibodies included: ADP ribosylation factor factor-like protein 13 B (Arl13b) (#17711-1-AP, Proteintech), Cathepsin D (#ab75852, AbCam), GFAP (#MAB360, EMD Millipore), LAMP-1 (#AB_528127, DSHB), LC3 (NB100-2220, NOVUS), S100β (#287 004, Synaptic Systems). Then, slices were washed three times 5 min in PBS before incubating with secondary antibody diluted in PBS containing 0.1% tween X-20, 5% NGS, during 1h at RT. Secondary antibodies included: Goat anti-guinea pig IgG Alexa Fluor 546 (#A-11074, Invitrogen), Goat anti-mouse IgG Alexa Fluor 546 (#A-11003, ThermoFischer), Goat anti-rabbit IgG Alexa Fluor 594 (#A-11037, ThermoFisher), Goat anti-rabbit IgG Alexa Fluor 647 (#A-21245, ThermoFisher) and Goat anti-rat IgG Alexa Fluor 680 (#A-21096, ThermoFisher). Afterwards, slices were washed 3 times 10 min in PBS. Hoechst 33258 (ThermoFisher) nuclear staining was added to the first PBS wash after the last secondary antibody. Sections were mounted onto microscope slides with Mowio/DABCOl mounting media and left overnight in darkness. For double/triple immunohistochemistry, primary and secondary antibody labelling was done sequentially.

### Confocal and STED microscopy

Confocal images were captured with Leica TCS Sp8 confocal microscope, following the same acquisition parameters between experimental and technical replicates (pixel size 16 bits; z-step 0.34µm; Plan Apochromat CS2 63x oil NA: 1,4).

For super-resolution STED microscopy of primary cilia, anti-rabbit conjugated with Alexa-488 Plus (Invitrogen) was used as secondary antibody and nuclei stained with DRAQ5 or DRAQ7. A 595 nm laser was used for depletion of Alexa-488 Plus. STED images were acquired with a 100X objective (numerical aperture = 1.4) with oil immersion at a pixel size ∼20 nm.

### Image analysis

For image processing and immunoreactivity quantification, NIH Image J software and Leica Application Suite X (LAS X) were used. Astrocyte morphological analysis was performed using the Simple Neurite Tracer plugin on Fiji following Tavares and colleagues’ instructions [15]. Percentage of ciliated astrocytes was calculated quantifying the number of PC (Arl13b+) presence over the total number of astrocytes (S100β+). Cilia length and width were quantified using the measure tool on both software. For puncta and puncta area quantification, a custom script was developed and modified by Federico N. Soria [16].

### Statistical analysis

Statistical analysis was carried out with GraphPad Prism 8.0.2. Two-way ANOVA test and Sidak’s multiple comparisons test were performed, both with 95% confidence interval. P values were represented by asterisks as follows: (*) p-value<0.05; (**) p-value >0.01; (***) p-value<0.001; (****) p-value<0.0001. Differences were considered significant when p-value<0.05.

## RESULTS

For this study we have characterized astrocytes as well as astrocytic cilia and autophagy in young (3mo) and old (15mo) Ift88-/-^Thy1^ and F/F mice. In these mice, the loss of neuronal cilia, as well as neuronal autophagy have been characterized previously and are the basis for the current study [14].

### Neuronal cilia loss reinforces age-dependent astrocyte reactivity in the hippocampus

To investigate whether astrocytes respond to the loss of PC in neurons (nPC), we started characterizing astrocyte GFAP changes in hippocampal CA1 from Ift88-/-^Thy1^ mice and their age matched F/F controls. Immunoblot analysis in hippocampal extracts (**Figure 1A**) showed increased GFAP levels in old mice, and in young Ift88-/-^Thy1^ mice (**Figure 1B**).

**Figure 1.**
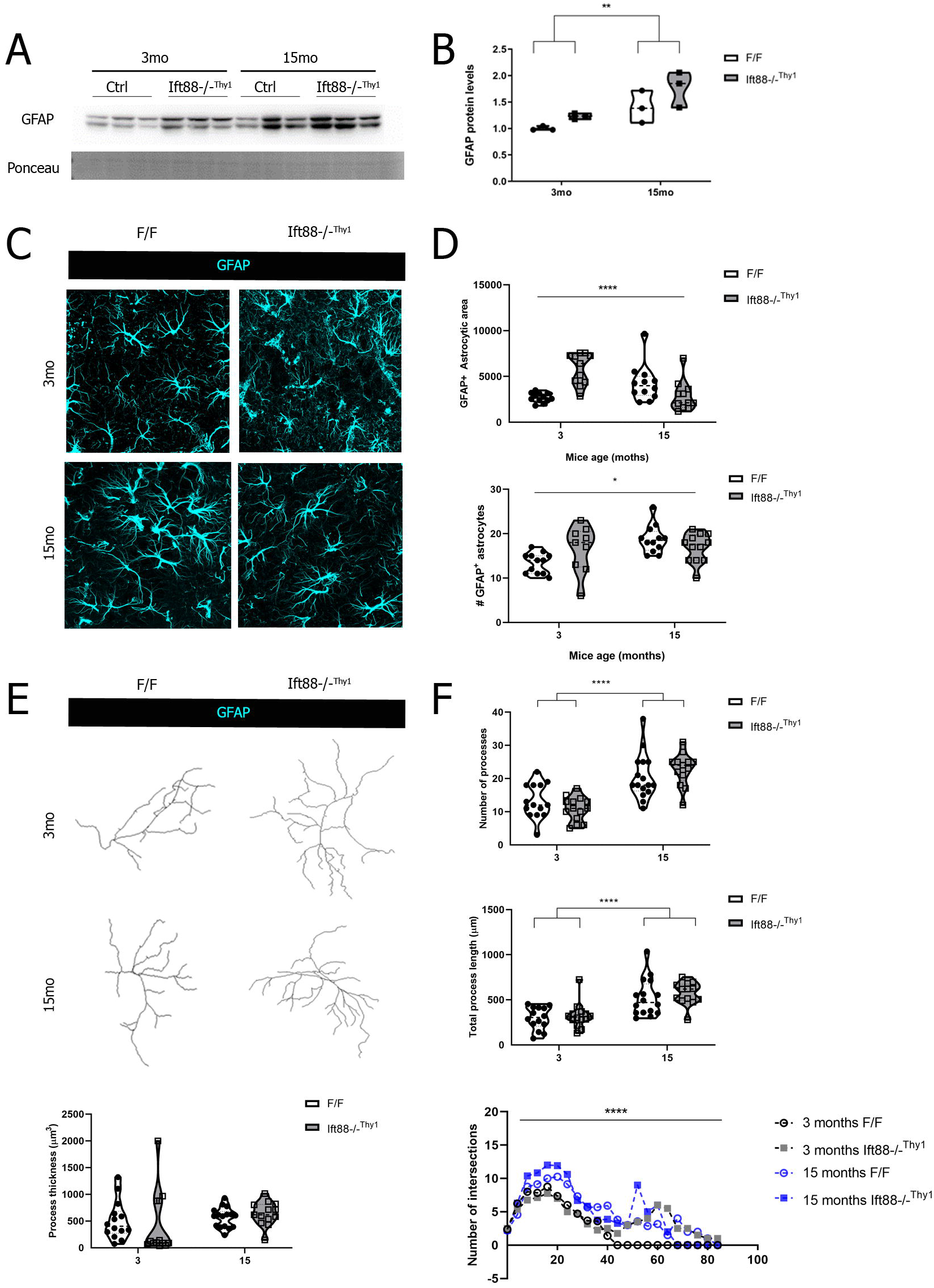
Astrocytes are reactive in Ift88-/-^Thy1^ mice. (A) Immunoblot against GFAP in hippocampal extracts from 3mo (young) and 15mo (old), F/F and Ift88-/-^Thy1^ mice. (B) Quantification of GFAP proteins levels from A. (C) Tissue immunofluorescence against GFAP (cyan) and confocal imaging in 3mo (young) and 15mo (old), F/F and Ift88-/-^Thy1^ mice. DAPI (grey) stains nuclei. (D) Quantification of the area occupied by GFAP (top) and the number of GFAP+ astrocytes (down) in C. (E) GFAP morphology analysis in 3mo (young) and 15mo (old), F/F and Ift88-/-^Thy1^ mice. (F) Quantification of the number of processes, process length and thickness, and Sholl analysis of E.

We then studied GFAP by tissue immunohistochemistry (IHC) in hippocampal CA1 (**Figure 1C**). Overall, area covered by GFAP staining and the number of GFAP^+^ cells increased in old mice, as well as in young Ift88-/-^Thy1^ mice (**Figure 1D**). We then assessed GFAP^+^ astrocytic structure as previously described (**Figure 1E**) [17]. Astrocyte morphometric analysis showed that aging increases the number of astrocytic processes, as well as their length, independently of the loss of nPC (**Figure 1F**). We did not observe changes in astrocytic process thickness. However, Sholl analysis showed that astrocyte morphology complexity increased (more process intersections and nuclei-distant branches) upon loss of nPC and was reinforced in old animals (**Figure 1F, down right**).

Altogether, GFAP analysis in CA1 from Ift88-/-^Thy1^ mice showed that the loss of nPC in young animals increases the protein levels, the area, the number, and the morphological complexity of GFAP^+^ astrocytes. Similar changes are observed in astrocytes from old F/F mice, which in addition have more and longer processes, and the loss of nPC in old animals boosts astrocyte morphology to a more reactive state.

### Ciliated astrocytes are reduced in old hippocampus

We next characterized the presence and morphology of primary cilia in CA1 astrocytes from young and old Ift88-/-^Thy1^ and F/F mice. As the PC is present in postmitotic, non-proliferating cells, we co-stained brain slices with Arl13b, a marker of astrocytic PC [18], together with the mature astrocyte marker S100β [19](**Figure 2A**). S100β stains preferably the soma of mature astrocytes and, considering that the PC is found close to the nucleus in the cell body, the use of somatic astrocyte marker eases to match Arl13b^+^ signal to astrocytes. In fact, similar to previous reports [18], our results show that Arl13b^+^ PC were present exclusively in S100β^+^ astrocytes (**Figure 2A, B down**), thus none Arl13b^+^ PC was found in other cell types such as EGFP-Thy1^+^ neurons (**Figure 2A down)**. Quantification in CA1 showed that the percentage of ciliated astrocytes decreased in old mice independently of the loss of nPC (**Figure 2B**) up, although we did no find changes in the length of this organelle (**Figure 2B middle**). We next took advantage of STED super-resolution microscopy to analyze the ultrastructure of astrocytic Arl13^+^ PC (**Figure 2C**). Quantification of cilia width at the tip and the base showed that astrocytic cilia width decreased in Ift88-/-^Thy1^ mice, and that aging abolishes this effect (**Figure 2D**).

**Figure 2.**
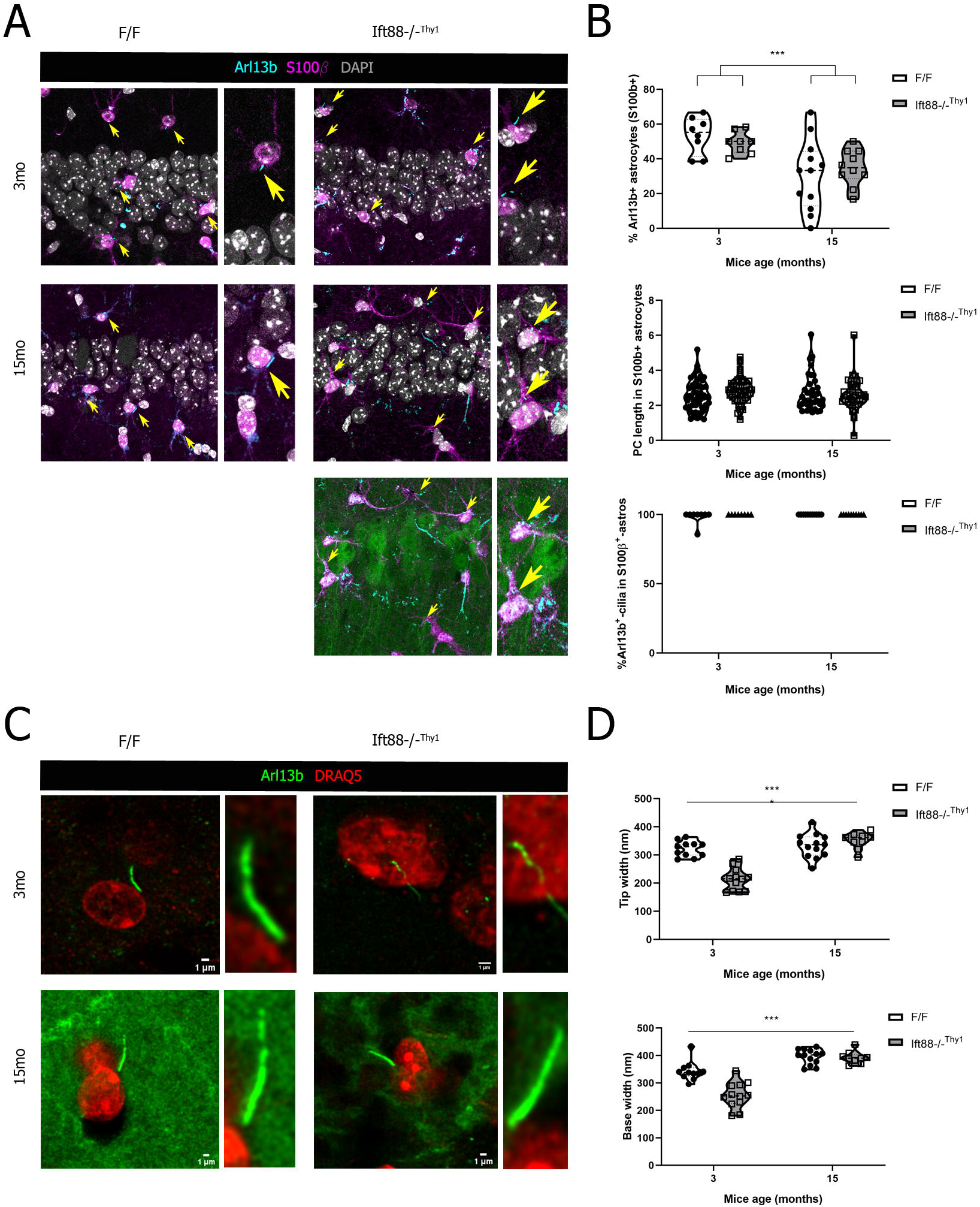
Astrocytic cilia changes in Ift88-/-^Thy1^ mice. (A) Tissue immunofluorescence against Arl13b (cyan) and S100β (magenta), and confocal imaging in 3mo (young) and 15mo (old), F/F and Ift88-/-^Thy1^ mice. (B) Quantification of the number of Arl13b+ ciliated astrocytes, length of Arl13b+ cilia, and percentage of Arl13b+ cilia in S100β+ astrocytes from A. (C) Tissue immunofluorescence against Arl13b (green) and nuclei staining with DRAQ5 (red), and STED superresolution imaging in 3mo (young) and 15mo (old), F/F and Ift88-/-^Thy1^ mice. (D) Quantification of Arl13b+ cilia width in C.

Overall we have found that, in contrast to neurons [14], astrocytes from old hippocampus are less ciliated independently of the presence of PC in neurons, a feature likely linked to the reactivity and thus, potential proliferative state of these cells in the aging brain. Additionally, loss of nPC elicits changes in the morphology of astrocytic cilia, which are reverted by aging.

### Autophagy in astrocytes responds to the loss of neuronal cilia

We have previously shown that signaling pathways in the PC modulate autophagy [13] and that in the hippocampus, neuronal ciliary pathways control autophagy in an age-dependent manner [14]. Here we have characterized by tissue immunofluorescence autophagy in CA1 astrocytes from young and old Ift88-/-^Thy1^ and F/F mice assessing LC3 and LAMP-1 (autophagosome and lysosomal marker, respectively) (**Figure 3A**).

**Figure 3.**
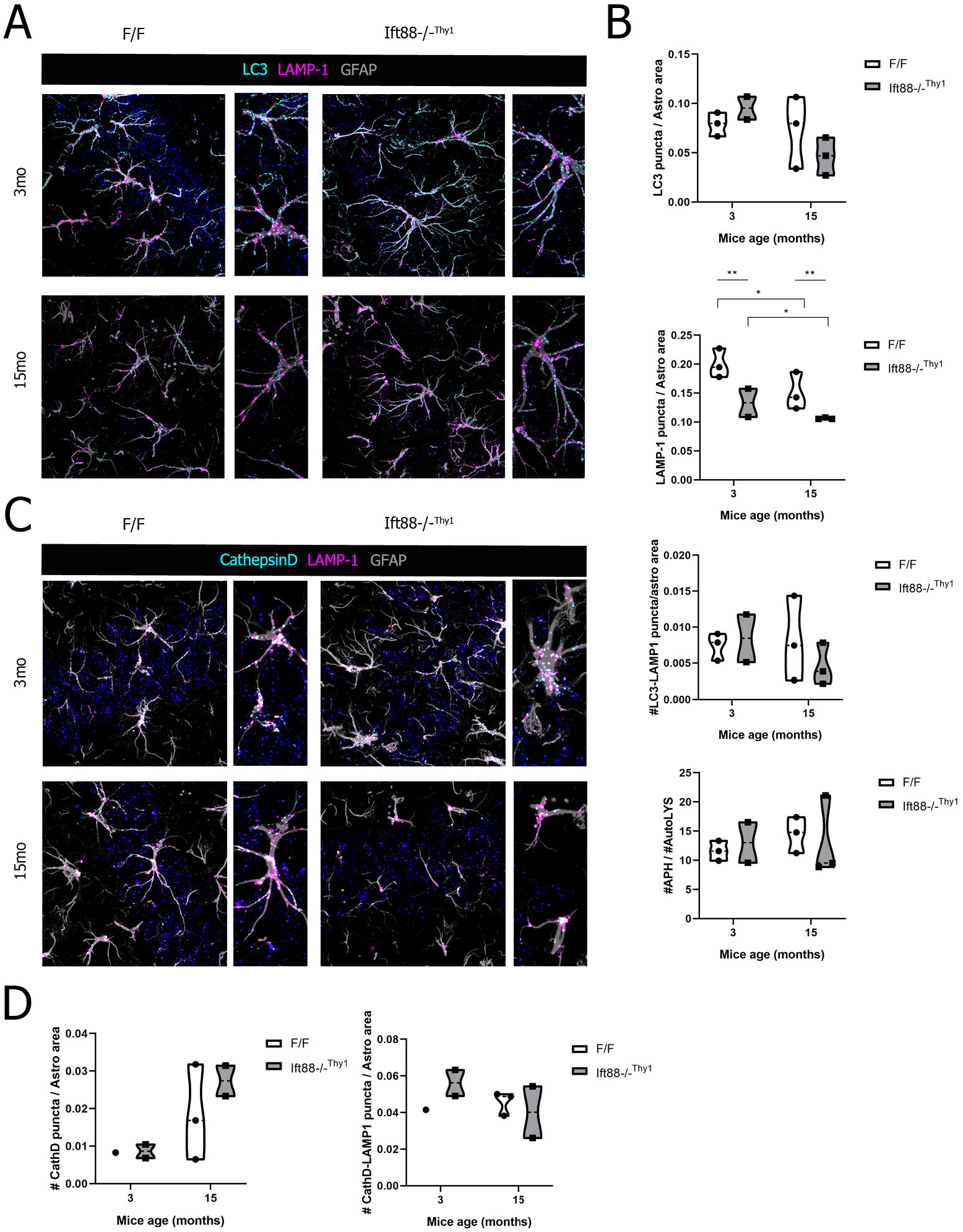
Astrocytic autophagy in Ift88-/-^Thy1^ mice. (A) Tissue immunofluorescence against LC3 (cyan), LAMP-1 (magenta) and GFAP (grey), and confocal imaging in 3mo (young) and 15mo (old), F/F and Ift88-/-^Thy1^ mice. (B) Quantification of the number of LC3+ puncta, number of LAMP-1+ puncta, number of LC3-LAMP-1 colocalized puncta, as well as the ratio between autophagosomes (APH) and autolysosomes (AutoLYS) in A. (C) (A) Tissue immunofluorescence against Cathepsin D (cyan), LAMP-1 (magenta) and GFAP (grey), and confocal imaging in 3mo (young) and 15mo (old), F/F and Ift88-/-^Thy1^ mice. (D) Quantification of the number of Cathepsin D+ puncta, and number of CathepsinD-LAMP-1 colocalized puncta in C.

LC3 vesicles did not show significant differences, but there is a trend towards lower LC3 puncta and area in old Ift88-/-^Thy1^ mice (**Figure 3B up**). In contrast, astrocyte LAMP-1 lysosomes decreased in Ift88-/-^Thy1^ mice independently of their age, and they were also diminished in the old animals independently of the genotype (**Figure 3B**). There was no change in the number of vesicles positive for both LC3 and LAMP-1 (**Figure 3B**) which are indicative of the fused autolysosomes. We also calculated the ratio between autophagosomes (LC3+ puncta) and autolysosomes (LC3+-LAMP-1+ puncta) as a proxy of autophagy flux in fixed tissue. Data show that there is no change in the calculated autophagy flux (**Figure 3B down**).

Since LAMP-1 was downregulated in astrocytes from mice lacking nPC, we assessed the status of lysosomal maturation by studying co-localization of Cathepsin D -a key enzyme in intralysosomal proteolysis [20]-with LAMP-1 lysosomes in GFAP^+^ astrocytes (**Figure 3C**). We found a trend to increasing number of Cathepsin D^+^ vesicles in old mice (**Figure 3D left**). However the number of LAMP-1 vesicles also positive for Cathepsin D in astrocytes did not change among experimental conditions (**Figure 3D right**), suggesting that the accumulation of Cathepsin D+ vesicles with aging occurs in non-LAMP-1 compartments. Considering that LAMP-1 is decreased in astrocytes from mice deficient in nPC with no change in the Cathepsin D content in these vesicles, we conclude that the overall degradation capacity of astrocyte lysosomes is decreased in young and old Ift88-/-^Thy1^ mice. Moreover, since maturation capacity of lysosomes is unaltered, we propose that the changes observed are likely due to a decrease in lysosomal formation.

## DISCUSSION

Astrocytes and neurons establish a bidirectional communication that helps maintaining brain homeostasis, a feature that is preserved during physiological aging [21]. In addition to cell-to-cell contact, astrocyte-neuron communication occurs through G-protein coupled receptors (GPCRs) [22] or exosomes [23]. As the PC is an organelle that senses extracellular changes and transduce them to modify intracellular events such as autophagy [13], here we hypothesize that astrocytes react to the loss of PC in neurons and modify their intracellular responses accordingly.

Our data show that neuronal cilia-depleted young mice acquire an astrocytic reactive phenotype. We have shown that GFAP protein levels and area occupied by GFAP+ astrocytes increase in young Ift88-/-^Thy1^ mice, together with the number of GFAP^+^ cells. Moreover, astrocyte morphology is more complex [24]. These results are in line to prior studies that have shown that loss of BBSome function, an adaptor protein complex that binds its cargo to the ciliary intraflagellar transport (IFT) complex [25], results in astrocyte activation [26].

We have also confirmed that, in old brains, astrocytes acquire a reactive phenotype [27; 28; 29] independently of the presence of PC. Furthermore, astrocytes from old Ift88-/-^Thy1^ mice had higher number and longer processes than young Ift88-/-^Thy1^ ones, and loss of neuronal PC boosted astrocyte morphological complexity. Altogether, these data indicate that loss of neuronal PC enhances aging driven astrocytic reactivity, increasing the deleterious effects of reactive astrocytes. Reactive astrocytes from aged brains have been linked to a pro-inflammatory phenotype [29], in contrast to reactive astrocytes from acute processes such as ischemia, which switch to a pro-survival phenotype [30]. Weather neuronal cilia loss associated astrocytic reactivity has a pro-inflammatory or pro-survival phenotype remains to be explored.

Our data also show that astrocytes from old brains are less ciliated, independently of the presence of PC in neurons. Reduced ciliation has been linked to increased proliferation in mesenchymal stem cells [31], a feature that might be also true in reactive astrocytes, also proliferative cells. Accelerated PC resorption might also result in decreased ciliation. In fact, neurons exposed to injury have a forced PC resorption that led to aberrant proliferation and ultimately apoptosis, and blockage of PC disassembly in these injured neurons rescues neuronal survival [32]. Thus, further research is needed to elucidate if, similar to injured neurons, reactive astrocytes accelerate cilia resorption and, consequently, increase proliferation

We also found that autophagy in astrocytes reacts to the lack of PC in neurons. For instance, neuronal cilia loss elicits a decrease in LAMP-1 astrocytic puncta, which is reinforced during aging. Autophagosomes in astrocytes show a trend to downregulation in old mice, reinforced upon loss of neuronal PC. Previously, we have shown that the absence of neuronal primary cilia promotes autophagy reductions in neurons [8]. Altogether, our data suggest that astrocytic autophagy changes might be a response to the decrease in autophagy in neurons or a direct consequence of the lack of neuronal PC.

Astrocyte-neuron interaction can occur through gliotransmitter release and clearance [7], calcium signaling [33], as well as through physical contact points, including integrin-Thy1 interaction at the plasma membrane [34]. In this interplay, Thy1 binds αvβ3 integrin and dephosphorylates p190Rho GTPase activating protein (GAP) [34]. Recent data have shown that point mutations in p190Rho GAP, which is located at the ciliary base, impairs ciliogenesis, and that cilia elongation requires a functional GAP domain [35]. Therefore, loss of PC in neurons could result in losing p190Rho GTPase activity that could in turn impact astrocyte-neuron communication. In addition, adjacent cells can contact each other by direct PC to PC contact, a type of cellular adhesion regulated by glycoproteins [36], or through extracellular vesicles (EV) excised from cilia [37; 38]. Astrocytes respond to EV through morphological changes in PC as well as by activating hedgehog signaling [39], a pathway that activates autophagy in a cilia-dependent manner [13]. Thus, the loss of neuronal PC might affect astrocytic autophagy, though further research is required to test this hypothesis.

The presented findings describe that astrocytes sense neuronal PC loss, which impacts the number and morphology of astrocytes, as well as their autophagic-lysosomal response. Moreover, these responses are disturbed during aging. These data are the first steps towards understanding the role of the PC as an intermediary in the communication between astrocytes and neurons, although more research is needed to elucidate its mechanisms and significance in health, aging and neurodegenerative diseases.

## AUTHOR CONTRIBUTIONS

OP conceptualized and designed the study, and acquired the funding. LdlH-G and OP prepared the tissue, performed the experiments, acquired and analyzed the images and the data. LdlH-G and OP wrote the manuscript.

## ACKNOWLEDGMENTS

We would like to thank Federico N. Soria for his support in image quantification and macro designing.

